# Essential role of the Conserved Oligomeric Golgi complex in *Toxoplasma gondii*

**DOI:** 10.1101/2023.05.30.542952

**Authors:** Clem Marsilia, Mrinalini Batra, Irina D. Pokrovskaya, Chengqi Wang, Dale Chaput, Daria A. Naumova, Vladimir V. Lupashin, Elena S. Suvorova

## Abstract

Survival of the apicomplexan parasite *Toxoplasma gondii* depends on the proper functioning of many glycosylated proteins. Glycosylation is performed in the major membranous organelles ER and Golgi apparatus that constitute a significant portion of the intracellular secretory system. The secretory pathway is bidirectional: cargo is delivered to target organelles in the anterograde direction, while the retrograde flow maintains the membrane balance and proper localization of glycosylation machinery. Despite the vital role of the Golgi in parasite infectivity, little is known about its biogenesis in apicomplexan parasites. In this study we examined *T. gondii* Conserved Oligomeric Golgi (COG) complex and determined that, contrary to predictions, *T. gondii* expresses the entire eight-subunit complex and each complex subunit is essential for tachyzoite growth. Deprivation of the COG complex induces a pronounced effect on Golgi and ER membranes, which suggests the *T. gondii* COG complex has wider role in intracellular membrane trafficking. We demonstrated that besides its conservative role in protein glycosylation and retrograde intra-Golgi trafficking, the COG complex also interacted with anterograde and novel transport machinery. Furthermore, we identified coccidian-specific components of the Golgi transport system: TgUlp1 and TgGlp1. Protein structure and phylogenetic analyses revealed that TgUlp1 is an adaptation of the conservative Golgi tethering factor Uso1/p115, and together with Golgi-localized TgGlp1, TgUlp1 showed dominant interactions with the trafficking machinery that predicted to operate the endosome-to-Golgi recycling. Together, our study showed that *T. gondii* has expanded function of the conservative Golgi tethering COG complex and evolved additional regulators of the transport likely to serve parasite-specific secretory organelles.

## INTRODUCTION

Apicomplexan parasites rely heavily on the proper function of their secretory pathways, which are much more complex than that of their host. In addition to conserved intracellular compartments, such as the ER, Golgi, trans Golgi network (TGN), endosomes, and plasma membrane, apicomplexans have several parasite-specific organelles. The novel organelles, including rhoptries, micronemes, dense granules, and the inner membrane complex (IMC), play central roles in parasite egress, invasion, and virulence making them attractive targets for research [1, 2]. Previous studies have established a close link between the biogenesis of these organelles and Golgi function [1, 3]. Golgi is a conserved eukaryotic organelle that processes and modifies secreted proteins. It has been shown that *T. gondii* expresses a pleura of secreted proteins that contain carbohydrate modifications, which would require passing through the Golgi [4–7]. Major virulence factors TgMIC2, TgAMA1, ROP18, and TgCST1 are among these glycan-modified proteins. While such importance and abundance of glycosylation explains why coccidian parasites retained a nearly complete Golgi system, most apicomplexan parasites reduced this compartment [8, 9]. Despite its significance, the role of the Golgi in apicomplexan secretory pathways remains an enigma.

Transport between membranous organelles is mediated by vesicular trafficking, which is executed in several steps. In the donor compartment, cargo is selected and packed into coated membrane vesicles that are recognized, docked, and fused with the acceptor compartment upon arrival. The process is highly specific for types of vesicles, donor and target membranes and involves a massive molecular machinery [10]. The major groups of regulators include cargo receptors, vesicle coat and its assembly factors, tethering machinery, membrane fusion SNARE receptors, and SNARE complex resolution factors. Selective capture, docking and fusion of the arriving vesicles is mediated by a versatile group of vesicle tethering factors, and specific SNARE molecules [11]. Bioinformatic analyses of apicomplexan genomes identified orthologs of several multi-subunit tethering complexes but could not detect long coiled-coil tethers [12, 13]. HOPS, COVET, and TRAPP tethering factors in *T. gondii* are localized in post-Golgi compartments, the trans-Golgi Network, and the endosome-like compartment (ELC), demonstrating their role in the biogenesis of secretory organelles [14–16]. However, Golgi tethering complexes are insufficiently studied.

A previous bioinformatic search identified four out of eight subunits of the conserved oligomeric Golgi (COG) complex in *T. gondii*, which is consistent with the current view of reduction in trafficking machinery across apicomplexan parasites [12]. The COG complex is a major Golgi multi-subunit tethering complex that has been extensively studied in model eukaryotes. The COG complex is organized in two lobes, lobe A and lobe B, and facilitates the tethering and fusion of Golgi recycling vesicles, which are responsible for retrieving escaped Golgi and ER resident glycosylation enzymes [17–19]. COG complex subunits interact with a selective set of trafficking machinery including COPI coat proteins, Golgi SNAREs, Rab effectors, and coiled coil tethers [20–27]. Acute depletion of individual or multiple COG complex subunits in mammalian cells led to massive accumulations of COG complex-dependent vesicles [28–30], while their complete knockout changed morphology of the Golgi and endosomal systems [31, 32]. It has been shown that malfunction in the COG complex alters N- and O-glycosylation of cellular proteins and causes several COG-related congenital disorders of glycosylation in humans [17, 33–35].

In the current study, we present experimental evidence that *T. gondii* expresses a complete oligomeric COG complex that is essential for tachyzoite growth, and we demonstrated its wider roles in parasites. We have found that the *T. gondii* COG complex interacts with novel Golgi coiled coil protein and identified a novel Golgi transport factor involved in retrograde transport in the late Golgi compartment.

## RESULTS

### *Toxoplasma* COG complex is composed of eight subunits

Phylogenetic analysis of the tethering complexes in metazoans identified four putative subunits of the octameric Golgi tethering COG complex in *T. gondii*, which could indicate that either the COG complex is reduced or the missing components were too dissimilar to be captured (Fig. 1) [12, 36]. To determine the exact composition of *Toxoplasma* COG complex, we examined proteins associated with TgCog8, whose counterpart in other eukaryotes functions as a bridge between lobe A and lobe B of the COG complex (Fig. 1B) [19, 37, 38]. The endogenous TgCog8 protein was fused with auxin-induced degron (AID) and 3xHA epitope tag (Fig. 1A) [39]. Co-staining with the Golgi marker TgGRASP55^RFP^ confirmed TgCog8^AID-HA^ was localized at the perinuclear Golgi (Fig. 1A). TgCog8^AID-HA^ protein complexes were isolated from dividing tachyzoites, and mass spectrometry followed by computational analysis detected 46 proteins with SAINT scores of 0.5 or higher (Fig. 1C and D, Table S1). Eight factors were particularly abundant: alongside TgCog8 we detected three predicted COG complex subunits, TgCog2, TgCog3, and TgCog4, and five proteins with unknown functions (Fig. 1D). We confirmed that four of these proteins were the missing COG complex subunits TgCog1, TgCog5, TgCog6, and TgCog7 (Fig. 1E, Fig. S1). TGME49_224150 contained a conserved Cog6 domain, suggesting that it is *Toxoplasma* Cog6 subunit. Phylogenetic analyses of TGME49_290310, TGME49_251730, and TGME49_242030 verified that these proteins were *Toxoplasma* counterparts of COG1, COG5, and COG7, respectively (Fig. S1). Further examination of amino acid sequences showed that TgCog5 contains a CATCHR (<u>c</u>omplexes <u>a</u>ssociated with tethering <u>c</u>ontaining <u>h</u>elical rods) fold (Fig. S2) [40]. Prior work in other eukaryotic models have shown that the binary COG5-COG7 subcomplex is stabilized by the antiparallel interactions of the α1-helix of the COG5 CATCHR fold and the N- terminal α-helixes of COG7 [41]. Coincidently, the sequence alignment and folding predictions revealed three α-helixes in the N-terminus of the TGME49_242030 protein, as well as amino acid residues that are critical for COG5-COG7 interaction, confirming the TGME49_242030 identity as TgCog7 subunit (Fig. S2). Together with phylogenetic evidence and the nearly equimolar levels of COG subunits detected in the TgCog8 proteome, our findings confirmed that *Toxoplasma* expresses a full complement of the eight subunits constituting the Golgi tethering COG complex.

**Figure 1.**
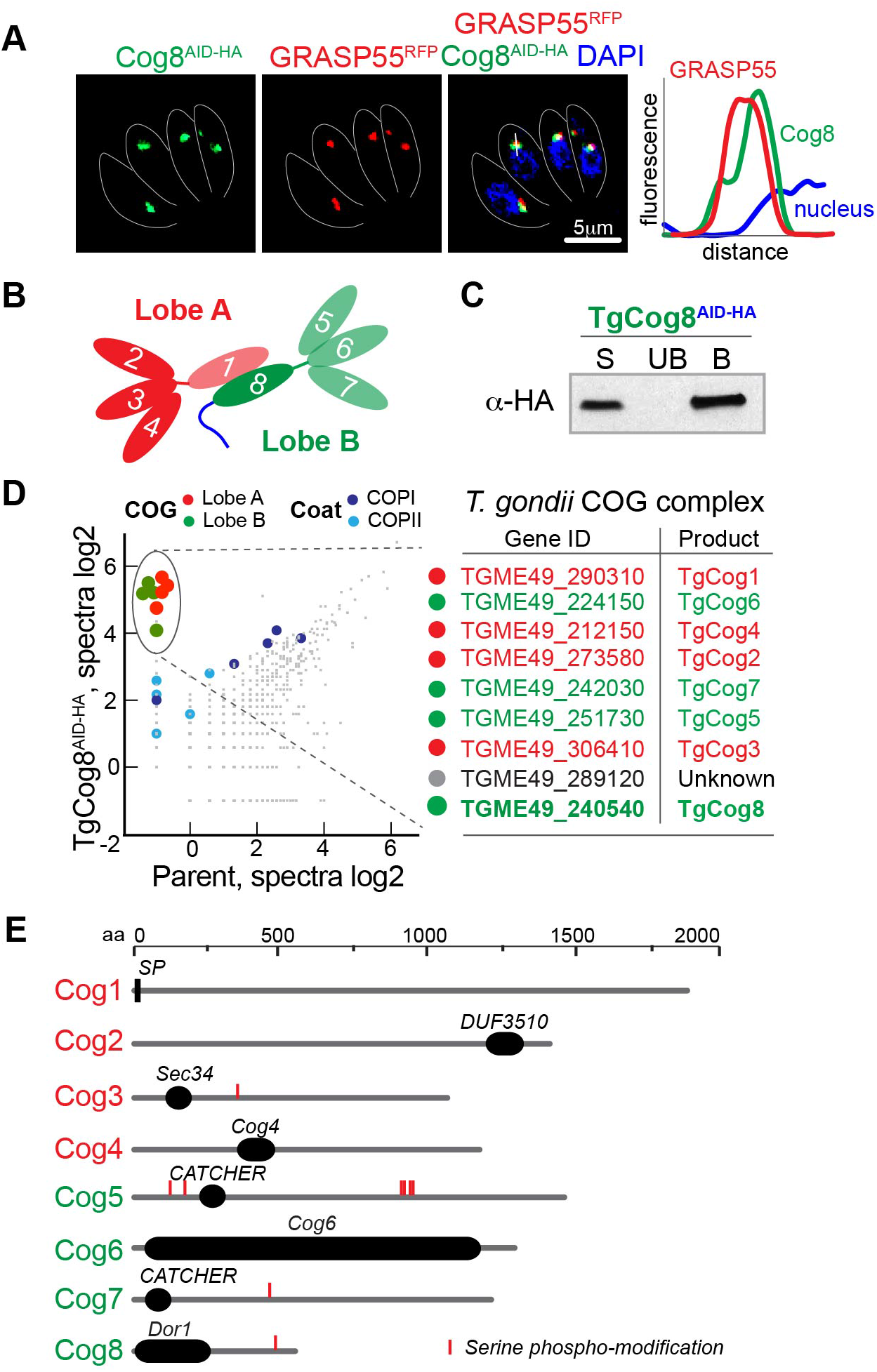
Identification of the *Toxoplasma* COG complex subunits. **A.** Endogenously tagged TgCog8^AID-HA^ (green) colocalizes with Golgi marker GRASP55^RFP^ (red). The blue DAPI stain marks tachyzoite nucleus. The intensity of TgCog8^AID-HA^, GRASP55^RFP^ and DAPI fluorescence in the region shown with the white bar is plotted on the graph to demonstrate significant colocalization of the COG complex subunit and the Golgi marker. **B.** The schematics shows two lobe composition of the COG complex. Previously identified subunits are shown with bright colors. **C.** Equal amount of the input fraction (S, soluble), proteins not retained on the αHA beads (UB, unbound) and proteins retained on αHA-beads (B, bound) were examined by western blot analysis to confirms efficiency of the TgCog8^AID-HA^ pulldown. **D.** The log2 values of the protein spectra detected by mass-spectrometry analysis of the TgCog8^AID-HA^ complexes are plotted on the graph. Maximally enriched proteins are encircled and listed in the table on the right. **E.** The diagram shows protein organization of the *Toxoplasma* COG complex subunits.

### *Toxoplasma* COG complex is essential for tachyzoite growth

To examine the function of COG complex in *T. gondii*, we generated conditional knockdown lines for each subunit. Subunits TgCog2 through TgCog8 were analyzed in the auxin-induced degradation model and TgCog1 was studied in the tet-OFF model [39, 42]. All the subunits of *Toxoplasma* COG complex were abundantly expressed in Golgi (Fig. 2A). We established that either treatment with auxin for 30 minutes (TgCog2 – TgCog8) or anhydrotetracycline for 16 hours (ATc) (TgCog1) both led to robust depletion of the target proteins (Fig. 2B). Contrary to their counterparts in host cells, each subunit of the *Toxoplasma* COG complex was found to be essential for tachyzoite survival. Persistent downregulation of any subunit was sufficient in preventing lytic plaque formation (Fig. 2C). However, downregulating individual subunits had various effects on parasite replication. While depleting of one of six subunits (TgCog2, TgCog4, TgCog5, TgCog6, TgCog7 and TgCog8) gradually arrested tachyzoite growth over multiple division cycles, depleting either TgCog1 and TgCog3 immediately affected parasite division (Fig. 2D). TgCog1- and TgCog3-deficient tachyzoites could not complete a second intravacuolar division cycle (an average 2 parasites per vacuole). These results demonstrate the vital role of the COG complex in the survival of *T. gondii* tachyzoites, specifically lobe A subunits TgCog1 and TgCog3.

**Figure 2.**
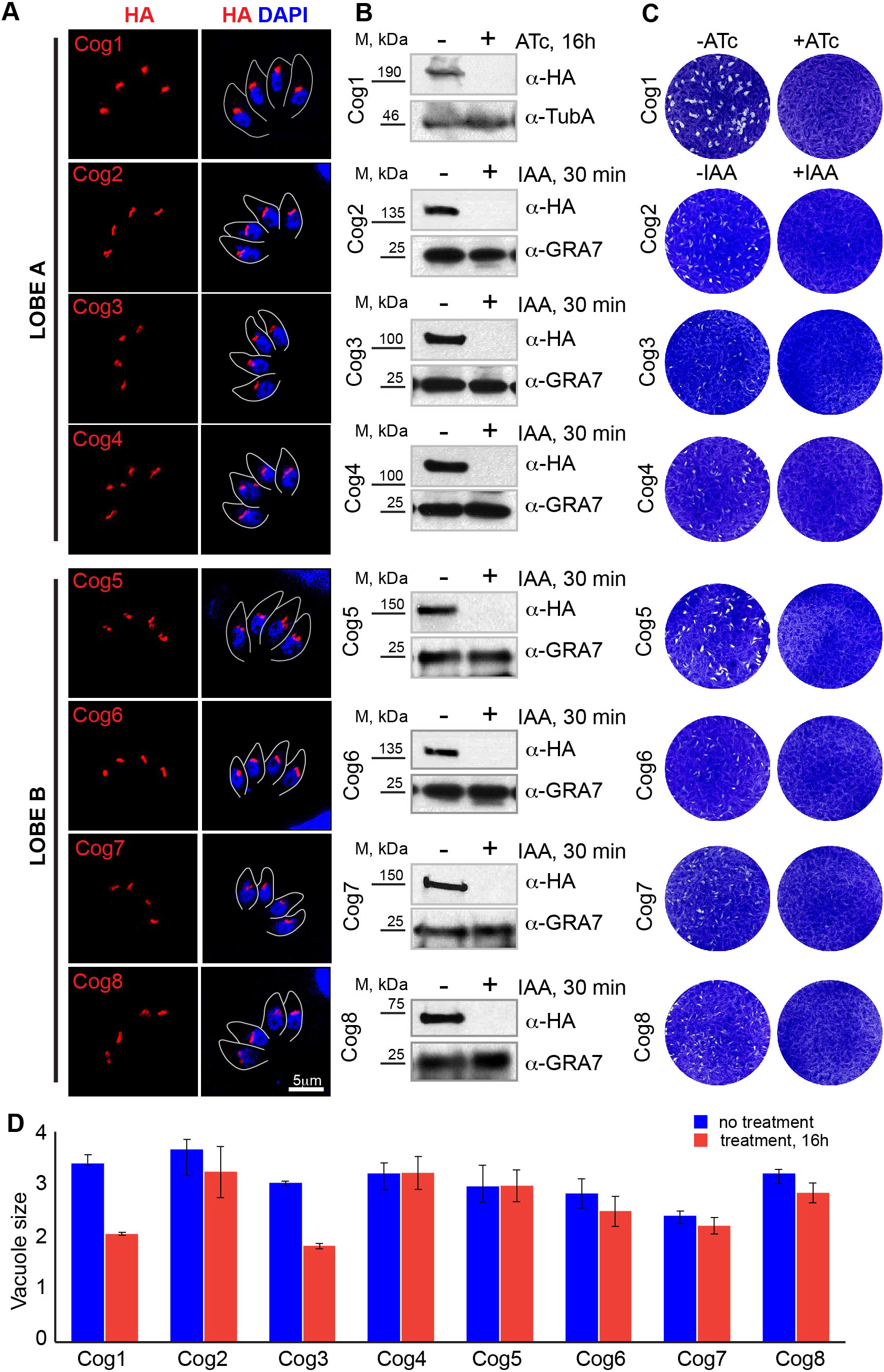
Eight essential subunits of the *Toxoplasma* COG complex. **A.** Immunofluorescent microscopy shows Golgi expression of the ^HA^TgCog1 tet-OFF model and AID models of TgCog2^AID-HA^ – TgCog8^AID-HA^ subunits using α-HA (red) and DAPI (blue) stain. The grey line represents parasite staining with antibodies against surface marker TgIMC1. **B.** Western Blot analysis confirmed expression and downregulation of ^HA^TgCog after 16-hour treatment with 2μM anhydrotetracycline (ATc) and TgCog2^AID-HA^ – TgCog8^AID-HA^ after 30-min treatment with 500μM auxin (indole-3-acetic acid, IAA). Western blots were probed with α-HA to detect the COG complex subunits, and either with α-Tubulin A or α-GRA7 to confirm equal loading of the total lysates. **C.** Images of the host cell monolayers infected with ^HA^TgCog1 tet-OFF or TgCog2^AID-HA^ – TgCog8^AID-HA^ AID tachyzoites and grown with or without indicated treatment for 7 days. **D.** The average number of parasites per vacuole after 16 hours growth in the presence or absence of 2μM ATc (TgCog1) or 500μM IAA (TgCog2 – TgCog8) was quantified in three independent experiments. Mean value -/+ SD are plotted on the graph.

### Rearrangement of *Toxoplasma* Golgi during cell division

To determine the function of the COG complex in *T. gondii*, we first reconstructed Golgi dynamics in dividing tachyzoites [9]. In eukaryotes, the Golgi expands and duplicates in synchrony with cell division to ensure it is inherited by both daughter cells [43]. We monitored Golgi morphology in tachyzoites by co-staining the COG complex subunit TgCog2^HA^ with budding stage marker TgIMC1 to visualize the assembly of internal daughters and centrosome marker TgCentrin1 to help to identify parasites in G_1_ cell cycle phase (Fig. 3). Our results showed that *Toxoplasma* Golgi undergoes significant rearrangement as the parasite progresses through endodyogenic cell division. *Toxoplasma* Golgi exists as a single compact organelle during G_1_, duplicates in S phase, breaks apart, and then restores its compact morphology during the internal budding process (Fig. 3A). We confirmed that *Toxoplasma* Golgi loses connection with the centrosome prior to centrosome duplication at the G_1_/S transition, where the centrosome translocates to the posterior end of the parasite nucleus (Fig. 3B, G_1_ and S panels) [44, 45]. The duplicated Golgi continues to grow and reconnects with the duplicated centrosomes, and each Golgi-centrosome unit is then delivered into the newly formed internal buds (Fig. 3A and B, budding progression). Golgi fragmentation is transient and characteristic to early budding (Fig. 3A, third panel), and is fully reassembled by mid-budding as a perinuclear rod-like structure at the apical end of the growing bud (Fig. 3A, bottom panel). We also examined the internal membranes of tachyzoites by transmission electron microscopy (TEM) and confirmed the compound *T. gondii* Golgi organization. In tachyzoites, the Golgi system is composed of 3-5 developed cisterns and a number of Golgi-associated vesicles. TEM analyses further verified that the Golgi is associated with apicoplast rather than the centrosome during the G_1_/S transition. *T. gondii* would need a repertoire of sophisticated regulatory machinery to maintain such complex organization of this essential and dynamic organelle, including the eight-subunit Golgi tethering COG complex.

**Figure 3.**
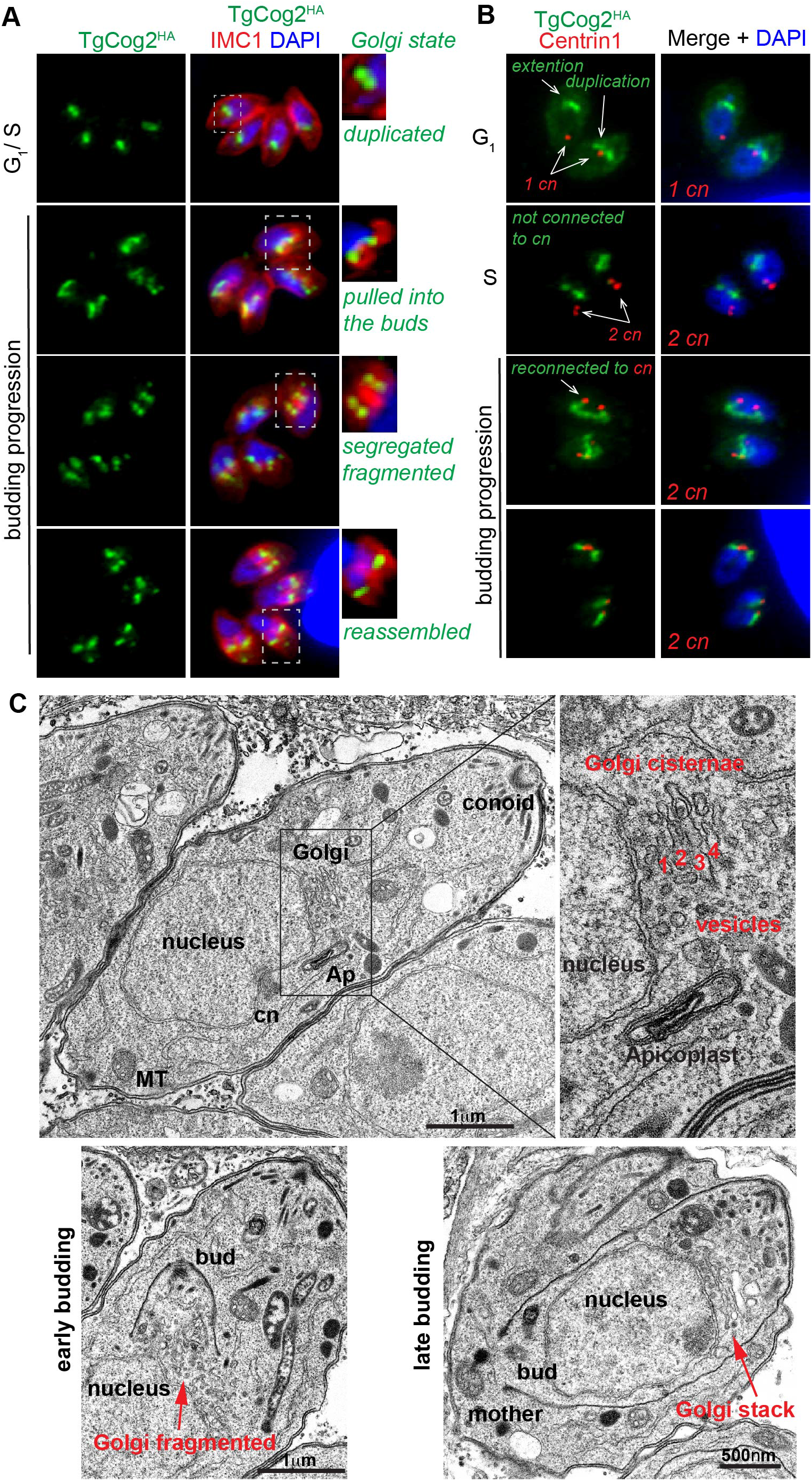
Cell cycle dynamics of the *Toxoplasma* Golgi. **A.** Major cell cycle phases were determined based on parasite cytoskeletal morphology (α-IMC1, internal budding) and shape of the nucleus (DAPI). Various Golgi states and transitions, including organelle duplication, segregation, fragmentation and reassembly were visualized using TgCog2^HA^ marker. **B.** Golgi-centrosome relationship was examined by co-staining TgCog2^HA^ and centrosome marker Centrin 1 (α-Centrin1). **C.** The TEM images of dividing tachyzoites. Major organelles and structures are labeled.

### *Toxoplasma* COG complex deficiency severely affects Golgi structure

Studies of the COG complex in mammalian cells revealed that siRNA and degron-assisted depletion of either COG3, COG4, or COG7 subunits dramatically changes Golgi morphology [28, 30, 46]. Recent TEM analyses of COG-deficient hTERT-RP1 and HeLa cells showed that uncoated COG complex dependent (CCD) vesicles accumulated and led to Golgi inflation, while prolonged COG3 deficiency caused severe Golgi fragmentation [28–30]. To evaluate the effects of COG complex deficiency in *T. gondii*, we examined TgCog3 AID (lobe A subunit) and TgCog7 AID (Lobe B subunit) tachyzoites by high-pressure freezing/freeze substitution TEM. In untreated tachyzoite controls, the Golgi was organized as a multi-cisternae stack. However, this defined Golgi structure was lost after a short 30-minute auxin treatment of TgCog3 AID parasites (Fig. 4A and B). The Golgi cisternae appeared disorganized, fragmented, and the Golgi region was packed with free-floating large vesicles (Fig. 4B). Prolonged TgCog3 deprivation resulted in complete restructuring of internal membranes (Fig. 4E); in addition to vesiculation, the largest membranous structure of the cell, ER, inflated and engulfed large portions of cytoplasm. Depleting the lobe B subunit TgCog7 had a partial effect on the Golgi system. Both short (30 minute) and long (8 hour) auxin treatments caused Golgi vesicles to accumulate but did not significantly affect the Golgi cisternae (Fig. 4C and D). We also did not detect the inflation of internal membranes, even after depleting TgCog7 for 8 hours. Our results suggest that the two lobes of the *Toxoplasma* COG complex, or certain subunits within the complex, provide different functions. This is consistent with findings in yeast, flies, and selected mammalian cell lines. The phenotypes we observed in COG complex-deficient *Toxoplasma* share a signature of COG deficiency with human cells: accumulation of Golgi vesicles upon the acute depletion of lobe A. Like in yeast and HeLa cells, knocking down TgCog3 has a stronger effect on Golgi structure than does downregulating COG7 [28]. However, contrary to studied eukaryotes, prolonged TgCog3 deprivation produced the inflation of an upstream secretory pathway compartment, the ER, rather than the resident compartment, Golgi, suggesting major differences in the pathways regulated by Toxoplasma COG and COG complex of host cells.

**Figure 4.**
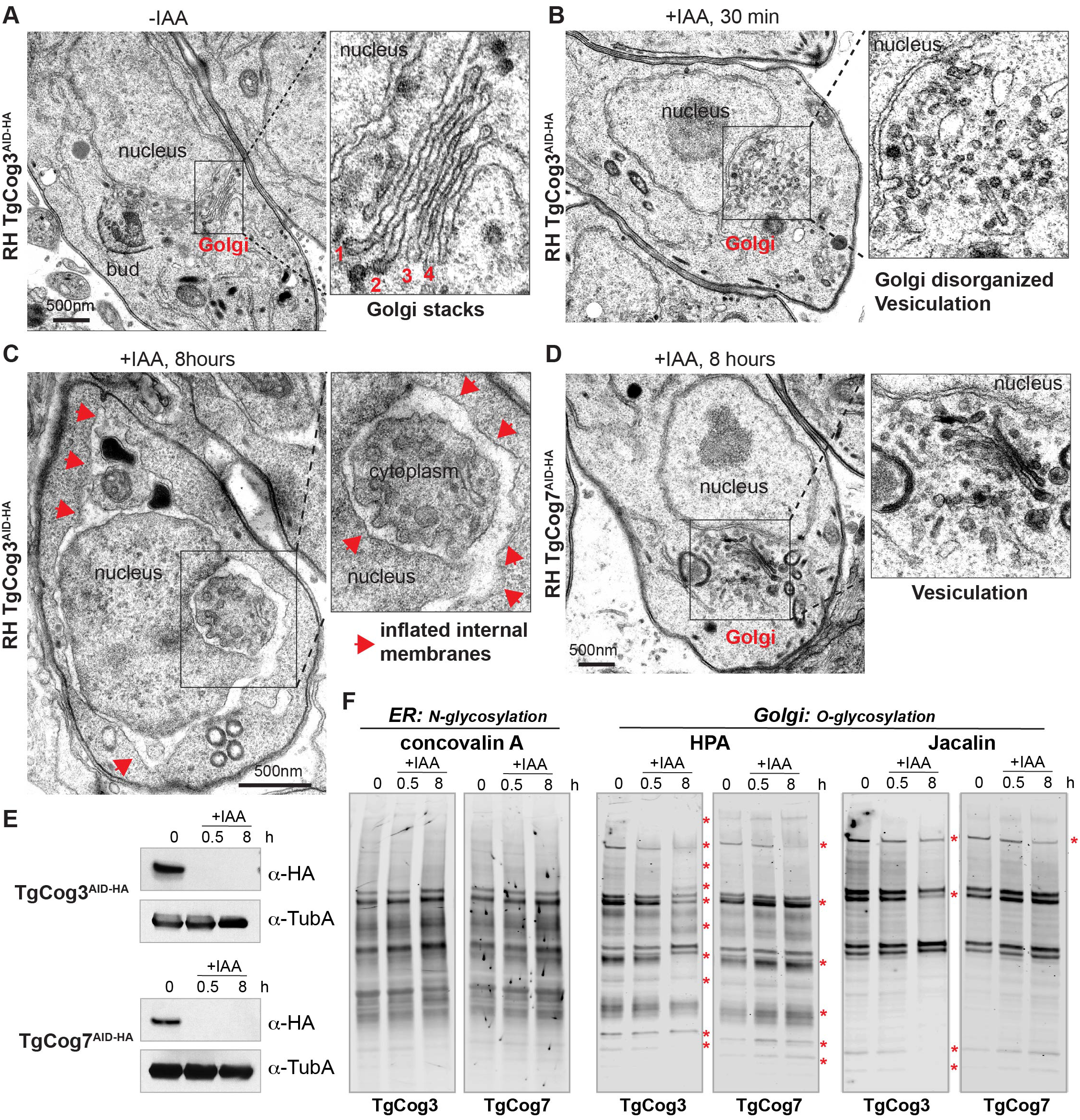
The COG complex is required for proper Golgi morphology and function. **A.** A TEM microphotograph of the tachyzoite expressing entire COG complex (-IAA). The enlarged image on the right depicts 4 cisterns of the Golgi apparatus. **B.** The image shows changes in the Golgi region of the RH TgCog3^AID-HA^ tachyzoite after 30 min IAA treatment. Note accumulation of the large Golgi vesicles. **C.** The prolonged TgCog3 deprivation on the tachyzoite internal membranes leads to inflation of the internal membranes, including ER (red arrowheads). **D.** The images depicted Golgi vesiculation in the RH TgCog7^AID-HA^ tachyzoite after 8 hours IAA treatment. **E.** Western Blot analysis confirmed efficient downregulation of TgCog3AID-HA and TgCog7AID HA after 30 min and 8 hours of IAA treatment. Western blots were probed with α-HA to detect the COG complex subunits, and with α-Tubulin A to verify equal loading of the total lysates. **F.** Images of the lectin-binding analysis. Total lysates of the IAA treated and not treated tachyzoites were probed with Concovalin A, HPA or Jacalin. Detected major changes are indicated with red stars.

### *Toxoplasma* COG complex deficiency alters protein O-glycosylation

Protein glycosylation is one of the most common post-translational modifications of secreted proteins, carried out by an extensive array of enzymatic machinery in the ER and Golgi to facilitate covalent attachment of oligosaccharides to specific amino acid residues. The elaborate process of glycosylation is executed in a precise order by specialized glycosyltransferases and glycosidases. Co-translational N-glycosylation of asparagine is carried out by ER enzymes, while O-glycosylation of serine or threonine side chains is catalyzed by Golgi-localized N acetylgalactosamine transferases [47]. Intracellular protein transport plays a central role in protein glycosylation, particularly the retrograde branch that restores the supply of glycosylation enzymes depleted by the continuous flow of anterograde transport. *T. gondii* encodes 68 glycogenes capable of assembling various types of glycans [5]. Studies using metabolic labeling alone or in combination with genome editing have demonstrated the proteins that contain the simple but abundant N- and O-glycosylations play important roles in parasite invasion, virulence, and environmental sensing [7, 48–50].

We determined that fragmentation and excessive vesiculation of the Golgi was the primary defect of downregulating TgCog3 and TgCog7. To find out whether these COG-induced morphological changes also affected Golgi function in tachyzoites, we used lectin staining to examine global protein glycosylation. Total lysates of tachyzoites expressing the complete COG complex (-IAA) or those depleted of TgCog3 or TgCog7 subunits for 30 minutes or 8 hours (+IAA) were probed with three types of glycan-binding lectins (Fig. 4E and F). The concanavalin A preferentially binds to α-mannose core chain of oligosaccharides present in N-glycans, while the HPA (*Helix pomatia* agglutinin) and Jacalin favor O-modified glycans. The selective binding of these lectins allows us to distinguish between ER- (N-glycosylation) and Golgi- (O-glycosylation) dependent modifications. Lectin staining revealed that tachyzoites lacking TgCog3 or TgCog7 had defects in Golgi-based O-glycosylation while ER-dependent N-glycosylation remained unaffected. Interestingly, although prolonged deprivation of TgCog3 leads to inflation of internal membranes, including the ER, this does not affect ER enzymes. This suggests that ER-based glycosylation machinery is not regulated by the COG complex. Consistent with our TEM studies, depleting TgCog3 has a more prominent effect on O-glycosylation than does downregulating TgCog7. Our glycosylation analyses corroborate the primary role of *Toxoplasma* COG complex in retrograde transport, particularly in recycling Golgi resident enzymes.

### *Toxoplasma* COG complex interacts with the coat proteins of retrograde and anterograde vesicles

The conventional role of the COG complex in eukaryotes is in tethering retrograde intra-Golgi vesicles [17, 33]. Two types of vesicles are used in ER-Golgi and intra-Golgi transport: anterograde COPII vesicles that deliver cargo from ER to Golgi are coated with four-subunit COPII complexes composed of Sec24, Sec25, Sec13 and Sec31 proteins, while the retrograde vesicles that restore balance of Golgi and ER enzymes and trafficking machinery are covered with seven subunit complexes containing α, β, β’, γ, δ, ε and ζ COPI proteins [51, 52]. The *Toxoplasma* orthologs of COPI and COPII complexes were previously identified *in silico* but neither anterograde nor retrograde Golgi transport had been studied in *T. gondii* (Fig. 5C and D) [53]. Analysis of TgCog8 pulldown identified equally enriched components of COPI and COPII coatomer complexes (Fig. 1, Table 1 and Table S4). To verify the interactions of *Toxoplasma* COG complex with both types of transport vesicles, we engineered transgenic lines co-expressing lobe A TgCog3^AID-HA^ or lobe B TgCog7^AID-HA^ proteins and subunits of COPI (TgCOPI-δ) or COPII (TgSec31) vesicles detected in the TgCog8 complexes. Immunofluorescent microscopy analyses confirmed Golgi localization of epitope tagged *Toxoplasma* coatomer proteins (Fig. S4A). We then examined the relationship between COG complex subunits TgCog3, TgCog7 and coat proteins by super-resolution microscopy and 3D reconstruction (Fig. 5A). Although both COPI and COPII vesicle coats had moderate to high probability of colocalizing with the COG complex, Pearson coefficient values suggested that *Toxoplasma* COG complex associates with retrograde COPI vesicles (TgCOPI-δ) more closely than with anterograde COPII vesicles (Fig. 5B). The strength of colocalization was comparable to that between COG complex subunits TgCog3 and TgCog7.

**Table 1.**
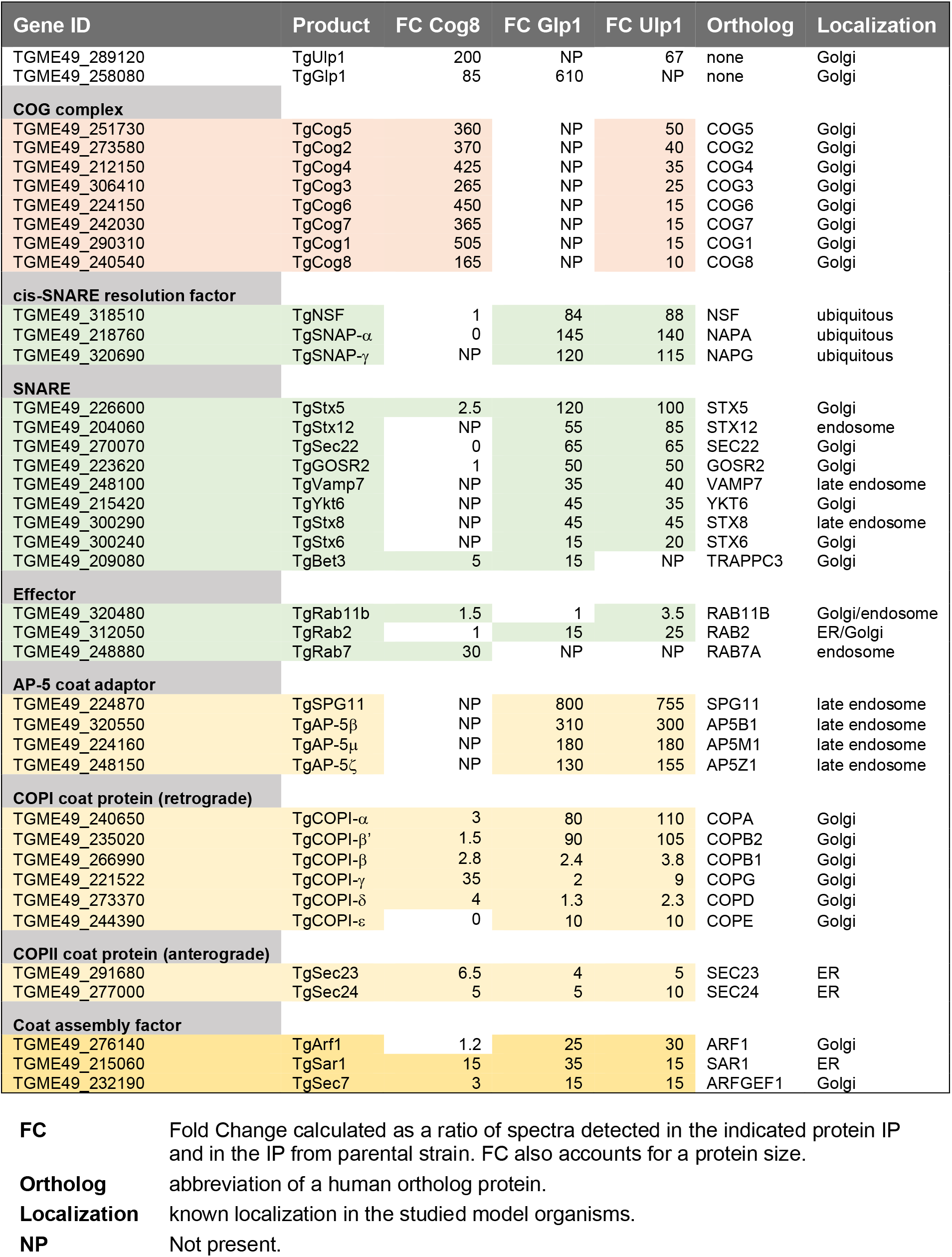
Summary of TgCog8, TgUlp1 and TgGlp1 protein interactions.

**Figure 5.**
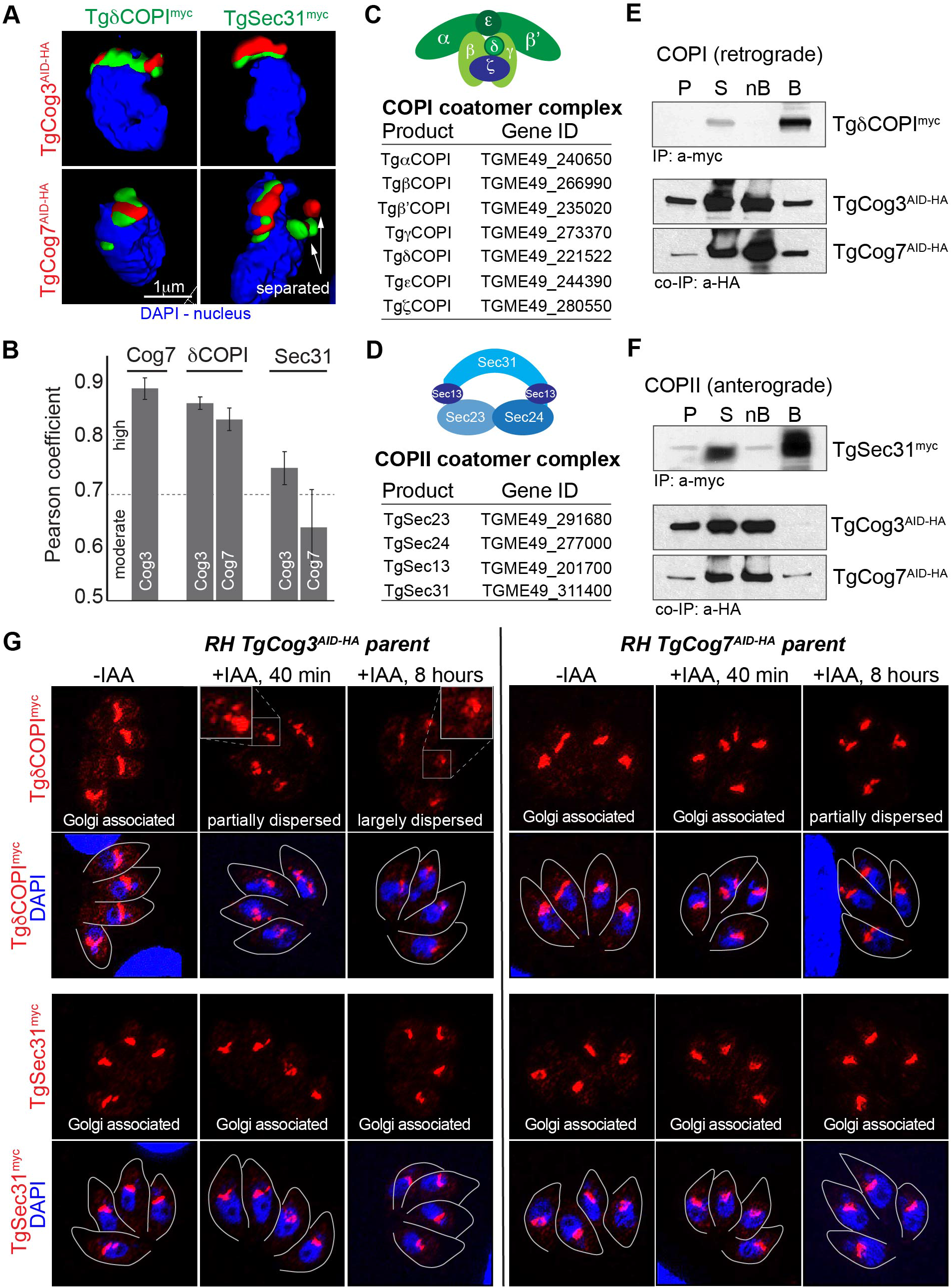
*Toxoplasma* COG complex interacts with COPI and COPII coatomer proteins. **A.** Images of the 3D reconstruction of the Golgi localized TgCog3^AID-HA^ (red), TgCog7^AID-HA^ (red), TgδCOPI^myc^ (green) and TgSec31^myc^ (green) in the perinuclear region (nucleus, DAPI, blue). **B.** Pearson coefficient was determined based on the colocalization analysis of a minimum of 10 tachyzoites. **C** and **D.** The diagrams of the COPI (C) and COPII (D) coatomer complexes and the tables of *T. gondii* orthologs of individual subunits. **E** and **F.** Immunoisolation of the TgδCOPI^myc^ (E) and TgSec31^myc^ (F) complexes from parasites co-expressing endogenous TgCog3^AID-HA^ or TgCog7^AID-HA^. The insoluble [P, pellet], soluble [S], depleted soluble fractions [nB, not bound] and the beads with precipitated complexes [B] (10 times more than the other fractions) were probed with α-myc and α-HA antibodies to detect potential interactions (the co-IP panels) and to confirm efficient pulldown of the bait protein (the IP panel). **G.** Immunofluorescent microscopy analysis of TgδCOPI^myc^ and TgSec31^myc^ coatomers in parasites expressing (-IAA) or lacking TgCog3^AID-HA^ or TgCog7^AID-HA^ (+IAA). Coatomers were visualized with α-myc antibodies and co-stained with DAPI (blue) and α-TgIMC1 (traced with grey line).

To verify the COG complex preference for retrograde transport, we performed co immunoprecipitation analysis of vesicle coat proteins and COG complex subunits (Fig. 5E and F). TgCOPI-δ^myc^ and TgSec31^myc^ complexes were affinity purified and probed for TgCog3^AID-HA^ or TgCog7^AID-HA^ proteins. Both COG complex subunits were detected in TgCOPI-δ^myc^ pulldown, and only TgCog7 in TgSec31^myc^ pulldown, verifying the COG complex interaction with retrograde COPI-coated Golgi vesicles. This suggests there exist specific interactions between the lobe B COG subcomplex and anterograde COPII-coated vesicles. However, relative abundance of co precipitated COG subunits suggests the *Toxoplasma* COG complex preferentially interacts with retrograde transport machinery.

We then tested how COG complex deficiency affects the expression of vesicle coat proteins. In line with co-immunoprecipitation results, we found the retrograde coatomer TgCOPI-δ^myc^ and not the anterograde coat subunit TgSec31^myc^ was substantially affected by TgCog3^AIDHA^ degradation (Fig. 5G). TgCOPI-δ^myc^ began to relocate to the cytoplasm after brief TgCog3^AIDHA^ absence (+IAA, 40 min), and by 8 hours, 100% of TgCog3 deficient tachyzoites had a pronounced loss of the Golgi associated TgCOPI-δ^myc^ (Fig. 5G and Fig. S4B). In addition, extended TgCog3^AID-HA^ deprivation (+IAA, 8 hours) led to a drastic decline of TgδCOPI^myc^ expression. Downregulating the lobe B subunit TgCog7^AID-HA^ had a lesser effect on TgCOPI-δ^myc^. Congruent with our TEM observations, the Golgi appeared unorganized due to TgCog7 deficiency (Fig. 4D), but TgCOPI-δ^myc^ remained associated with fragmented Golgi membranes in 50% of TgCog7-deprived tachyzoites. (Fig. S4B). These results further confirmed that the *Toxoplasma* COG complex predominantly interacts with retrograde transport machinery. However, contrary to studies in model eukaryotes, *Toxoplasma* COG complex also demonstrated involvement with anterograde transport, suggesting overlapping function in regulating both forward and recycling vesicles.

### *Toxoplasma* COG complex interacts with coccidian Uso1-like Golgi tethering factor

Like the other COG complex subunits, TgCog8 also strongly interacted with a coccidian-specific protein of unknown function TGME49_289120 (Fig. 1D). To confirm TGME49_289120 expression and interaction with *Toxoplasma* COG complex, we generated the TGME49_289120 AID conditional expression model in RH*ΔKu80Δhxgprt AtTIR1* parasites. Immunofluorescence microscopy analysis revealed its localization at the Golgi localization and a plaque assay showed that TGME49_289120 was essential for tachyzoite growth (Fig. 6A and B). We also created TgCog3 and TgCog7 AID models expressing epitope tagged TGME49_289120^myc^ and verified TGME49_289120 colocalization with COG complex (Fig. 6C and Fig. S4C). To test interaction with COG complex, we examined TGME49_289120^myc^ complexes by mass-spectrometry (Fig. 6E, Table S). Corroborating proteomic studies of TgCog8, TGME49_289120 protein complexes contained a complete set of COG complex subunits, and we confirmed the presence of TgCog3^AID-HA^ and TgCog7^AID-HA^ by Western blot analysis (Fig. 6D and Table 1). Finding a complete set of COG subunits indicated that TGME49_289120 interacts with the entire COG complex. Relative abundance of COG subunits suggested that TGME49_289120 interacted with COG complex via TgCog5, TgCog2 and TgCog4 subunits.

**Figure 6.**
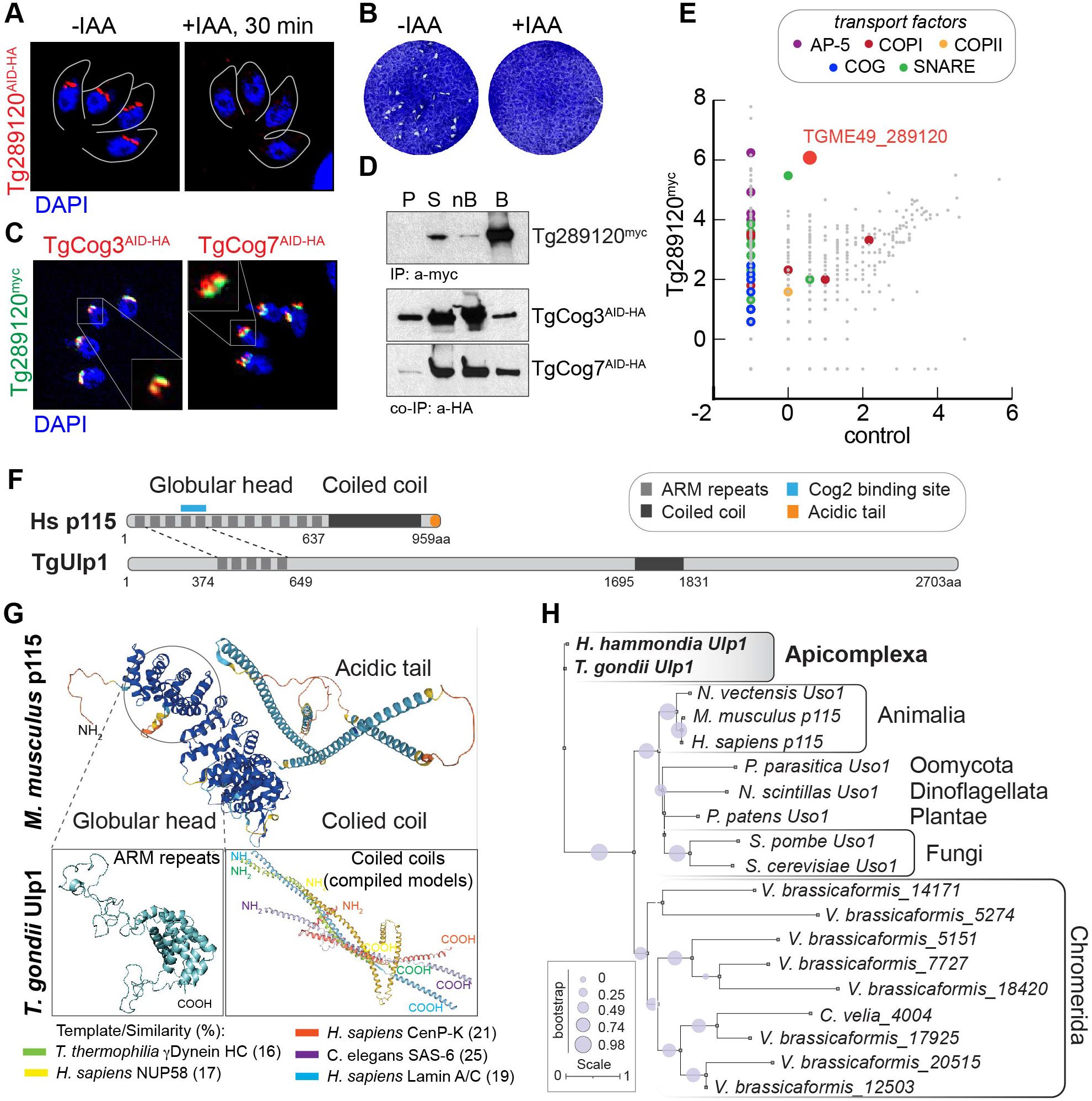
*Toxoplasma* COG complex interacts with novel Uso1-like factor. **A.** Localization of TGME49_289120^AID-HA^ protein was determined by co-staining of the factor (α HA), nuclear stain DAPI and parasite surface marker TgIMC1 (traced with grey line). A 30-min treatment with 500μM IAA resulted in robust TGME49_289120^AID-HA^ downregulation. **B.** Images of the host cell monolayers infected with Tg289120^AID-HA^ tachyzoites and grown with or without 500μM IAA for 7 days. **C.** Immunofluorescent analysis of the parasites expressing Tg289120^myc^ in the RH TgCog3^AID-HA^ or TgCog7^AID-HA^ mutants. Parasites were co-stained with α-myc (green), α-HA (red) and DAPI (blue). Insets shows the proteins overlap in the Golgi region. **D.** Western Blot analysis of immunoprecipitated Tg289120^myc^ complexes from parasites co-expressing endogenous TgCog3^AID-HA^ or TgCog7^AID-HA^. The insoluble [P, pellet], soluble [S], depleted soluble fractions [nB, not bound] and the beads with precipitated complexes [B] (10 times more than the other fractions) were probed with α-myc and α-HA antibodies to detect protein interactions (the co-IP panels) and to confirm efficient Tg289120^myc^ pulldown (the IP panel). **E.** The log2 values of the protein spectra detected by mass-spectrometry analysis of the Tg289120^myc^ complexes are plotted on the graph. The different color dots represent categories of the transport proteins. **F.** Schematics of *H. sapiens* p115 and *T. gondii* Ulp1 protein organization. The signature domains and regions of similarity are shown. **G.** Folding prediction for *M. musculus* p115 and selected regions of *T. gondii* Ulp1 are shown (AlphaFold2, PyMol, SwissProt). Note that image of the coiled coil region of *T. gondii* Ulp1 is a compilation of five different models listed in the legend below. **H.** Phylogenetic tree of TgUlp1 and various Uso1/p115 orthologs.

Detailed analyses of interactions strongly indicated that TGME49_289120 is involved in retrograde transport, further supported by its preferential interaction with Adaptor Protein complex 5 (AP-5), derived from late endosomes and Golgi COPI coat proteins (Table 1, Table S4). Both types of vesicles are implicated in retrograde protein transport [54, 55]. We had detected four components of AP-5/SPG11 complex and six COPI subunits with relative enrichment of outer layer COPI components (α and β’ subunits). In addition, orthologs of the major Golgi (TgStx5, TgGOSR2, TgSec22, TgYkt6, TgStx6) and endosome (TgStx12, TgVamp7, TgStx8) SNARE complexes, were among the strongest TGME49_289120 interactors [14, 56, 57]. Finally, a substantial presence of cis-SNARE complex resolution factors, TgNFS, TgSNAP-α, and TgSNAP-γ pointed toward a role for TGME49_289120 in vesicle tethering and fusion rather than in assembly and budding [10]. The enrichment of late endosomal and Golgi factors in Tg289120 complexes suggested that the role for this novel coccidian-specific factor lies in late endosome-to-Golgi and intra-Golgi transport.

Searching for clues of TGME49_289120 identity, we performed comprehensive analyses of protein folding. We found that TGME49_289120 protein organization was like that of the eukaryotic tethering factor p115/Uso1 (Fig. 6F). Uso1/p115 is a long coiled-coil protein that promotes the capture, docking, and fusion of vesicles traveling between ER and cis-Golgi [58]. Previous bioinformatic analyses suggested that apicomplexans lack conserved coiled-coil Golgi tethers, including Uso1 orthologs [12, 36]. Our examination of TGME49_289120 structure revealed that, while vastly different at the amino acid sequence level, TGME49_289120 contained several multi-helical armadillo-repeats (ARM) in the N-terminus predicted to fold in a structure reminiscent of the globular head domain of p115/Uso1 (Fig. 6G) [59]. Interestingly, this region has a substantial degree of amino acid residue similarity (21%) with human p115 and includes two ARM repeats that were previously shown to interact with COG complex via COG2 subunit [24]. Coincidently, TgCog2 was among the *Toxoplasma* COG subunits that preferentially interacts with TGME49_289120 (Table 1). Despite the lack of similarity, TGME49_289120 was also predicted to have a coiled-coil region in its C-terminus like Uso1/p115 (1696-1830) (Fig. 6F and G). Corroborating the prediction that a true ortholog was lost in evolution, our phylogenetic analyses determined that TGME49_289120 was not related to the eukaryotic Uso1/p115 tethering factors (Fig. 6F). The similarities in structure, localization at the Golgi, and predicted roles in tethering vesicles between TGME_289120 and Uso1/p115 suggested that coccidian parasites reinvented a novel Golgi tethering factor, and we named it TgUlp1 (<u>U</u>so1-like <u>p</u>rotein 1).

### *Toxoplasma* expresses a novel Golgi transport factor

Another coccidian-specific protein with unknown function TGME49_258080 was detected in the TgCog8 interactome (Table S). To gather clues on its function, we built the TGME49_258080 AID model and we found that TGME49_258080 was not essential for parasite replication and, similarly to the COG complex, TGME49_258080 was expressed on the tachyzoite Golgi (Fig. 7A and B). Super resolution microscopy analyses of TGME49_258080^myc^ co-expressed with either TgCog3^AID-HA^ or TgCog7^AID-HA^ confirmed that the proteins are highly likely to colocalize (Fig. 7C, Fig. S4D and E). Examination of TGME49_258080 protein folding revealed two coiled-coil regions in the C-terminus with reasonable similarity to those in proteins that are involved in intracellular protein transport in various eukaryotic models, including the conserved guanine nucleotide exchange factor Sec2 and intraflagellar transport protein from *Chlamydomonas reinhardii* IFTB1 (Fig. 7D, Fig. S4E) [60, 61]. We decided to name this novel Golgi localized TGME49_258080 protein TgGlp1 (<u>G</u>olgi *p*rotein 1).

**Figure 7.**
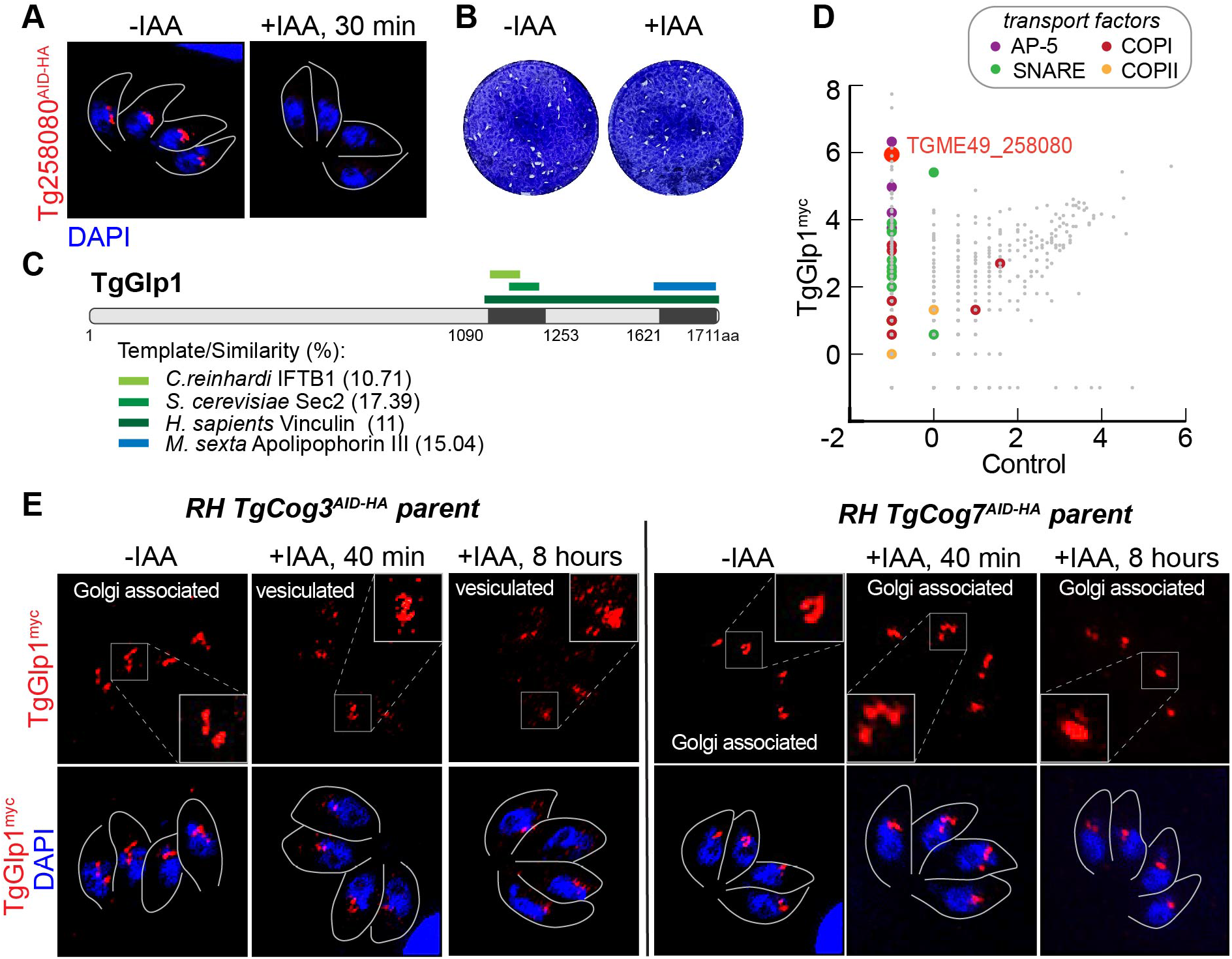
TgGlp1 is novel *T. gondii* Golgi transport factor. **A.** Localization of TGME49_258080^AID-HA^ protein was determined by co-staining of the factor (α HA), nuclear stain DAPI and parasite surface marker TgIMC1 (traced with grey line). A 30-min treatment with 500μM IAA resulted in robust TGME49_258080^AID-HA^ downregulation. **B.** Images of the host cell monolayers infected with Tg258080^AID-HA^ tachyzoites and grown with or without 500μM IAA for 7 days. **C.** Schematics of TgGlp1 protein organization. The identified domains are shown. The regions of similarity are listed below. **D.** The log2 values of the protein spectra detected by mass-spectrometry analysis of the TgGlp^myc^ complexes are plotted on the graph. The different color dots represent categories of the selected transport proteins. **E.** Images of the tachyzoites expressing or lacking TgCog3 or TgCog7 for indicated time. The associated changes in the TgGlp1^myc^ expression were visualized by co-staining parasites with α-myc and DAPI (blue). The insets are overexposed images of the selected Golgi regions.

Mass spectrometry analysis of isolated TgGlp1^myc^ complexes did not show *Toxoplasma* COG complex subunits, suggesting that interactions between TgGlp1 and COG complex are indirect (Fig. 7D, Fig. S4G, Table 1, Table S). Nevertheless, the TgGlp1^myc^ proteome had several hits in common with the TgCog8 interactome, indicating that the COPI coat was the best candidate for the link with COG complex. Despite the lack of direct interaction, we found that COG complex significantly affected TgGlp1 expression. Downregulation of TgCog3 and, to a lesser extent, TgCog7, led to a dramatic reduction and relocation of TgGlp1 from its resident compartment to dispersed associated punctata (Fig.7E and Fig. S4C). Since TgGlp1 did not directly interact with COG complex, the observed changes were likely due to TgGlp1 associating with the Golgi region that is highly responsive to acute degradation of TgCog3 or TgCog7 as demonstrated in our TEM experiments (Fig. 4). Interestingly, TgGlp1 complexes had substantial overlaps with TgUlp1 proteome, implying these two novel coccidian specific Golgi proteins have similar functions (Table 1). Like TgUlp1, TgGlp1 preferentially binds AP-5 and COPI coat proteins and select Golgi and late endosomal SNAREs, indicating their involvement in tethering and docking the recycling vesicles to Golgi. Enriched cis-SNARE factors further indicate a specific role for TgGlp1 and TgUlp1 in disassembling post-fusion SNARE complexes.

## DISCUSSION

Golgi is a complex membranous organelle that functions as a transit station. It accepts vesicles from ER (anterograde transport) and late endosomes (retrograde flow of endosomal system) and sends vesicles to late secretory compartments such as TGN, late endosomes, and plasma membrane (anterograde), and then back to ER (retrograde transport). In addition, Golgi constantly recycles its resident proteins in trans-to-cis fashion (retrograde transport). Therefore, according to the current cisternae maturation model, Golgi is equilibrated by the anterograde and retrograde flows of transport intermediates. These two directions of protein transport have different goals. Anterograde transport delivers ER and Golgi processed cargo to its destination, while retrograde transport recycles components of trafficking machinery and modifying enzymes to their places of function. How Golgi segregates these two flows is still under investigation. However, it has been shown that model eukaryotes employ distinctive molecular machinery; anterograde transport is mostly mediated by COPII, clathrin, and AP-4 coated vesicles, while retrograde transport largely uses COPI and AP-1, 2, 3, and 5- coated vesicles [62].

Most apicomplexans have rudimentary Golgi systems, which can explain the lack of research attention to this organelle. Previously published data, and our study, demonstrate that *T. gondii* tachyzoites have a well-developed Golgi system composed of multiple cisterns and associated vesicles, corroborating retention of Golgi ribbon formation factors GRASP65/GRASP55 in the parasite genome [9]. This observation sets coccidian parasites apart from the phylum and suggests an important role of Golgi-dependent processes in parasite survival. It also explains why *T. gondii* has preserved substantial transport machinery, while most apicomplexan parasites have lost many of these components [12, 36]. Bioinformatic and empirical analyses identified *T. gondii* orthologs of all known vesicle coats and fusion factors, including SNAREs, Rabs, and cis-SNARE resolution proteins [1, 12, 14, 36, 53, 56]. However, these studies predicted variable degrees of preservation of vesicle tethering machinery, with a lack of coiled coil tethers and a structurally reduced COG complex [12, 36].

Our study showed that *T. gondii* preserved a full set of subunits of the Golgi tethering COG complex. We demonstrated that *T. gondii* COG complex has conserved function and displays several novel features. Like its higher eukaryote counterpart, *T. gondii* COG is important for protein glycosylation and preferentially interacts with the COPI vesicular coat, which confirms the conventional COG role in retrograde transport [17, 33]. However, unlike in its eukaryotic counterparts, each complex subunit was essential for tachyzoite growth, raising the importance of Golgi tethering factors and the contribution of Golgi to parasite survival. Although the immediate effect of *T. gondii* COG complex downregulation resembled the massive release of Golgi associated vesicles observed in mammalian cells, the long-term effect was very different (Fig. 8). COG complex deprivation in HeLa, HEK293T, and RPE1 human cells led to the inflation and severe fragmentation of Golgi but it was still capable of supporting anterograde transport, while tachyzoites deficient in lobe A subunit TgCog3 had pronounced bloating in the ER that presented as a severely detached nuclear envelope [31, 63]. The likely scenario of the events that lead to ER inflation caused by COG insufficiency is that *T. gondii* Golgi would fail to accept anterograde ER-derived vesicles, blocking secretion. In human cells, COG depletion is partially compensated by a large group of COG-independent coiled-coil tethers (V.L. personal communication), while *T. gondii* lacks this compensatory mechanism. It is also possible that in COG-depleted cells, Golgi fails to accept recycling vesicles arriving from late secretary compartments, causing non-specific fusion with the largest cellular organelle, ER. The presence of COPII coats of the anterograde vesicles and SNAREs circulating between ER and Golgi (Stx5 and Bet3) in the TgCog8 interactome support that *T. gondii* COG complex is involved in anterograde transport. This observation contradicts findings in mammalian cells where the COG complex does not interact with anterograde machinery and thus does not regulate anterograde trafficking. Interestingly, investigation of the temperature-sensitive mutants in yeast indicated that yeast COG complex is involved in ER-Golgi anterograde trafficking [64, 65]. This suggests some similarities between COG function in yeast and *T. gondii*. The broad role of the COG complex in intracellular trafficking may explain the unique essentiality of this complex in *T. gondii*. Identification of the interactors of individual COG complex subunits and deciphering the composition of COG complex-dependent vesicles will shed light on their origin and expand our understanding of the COG complex function in *T. gondii*.

**Figure 8.**
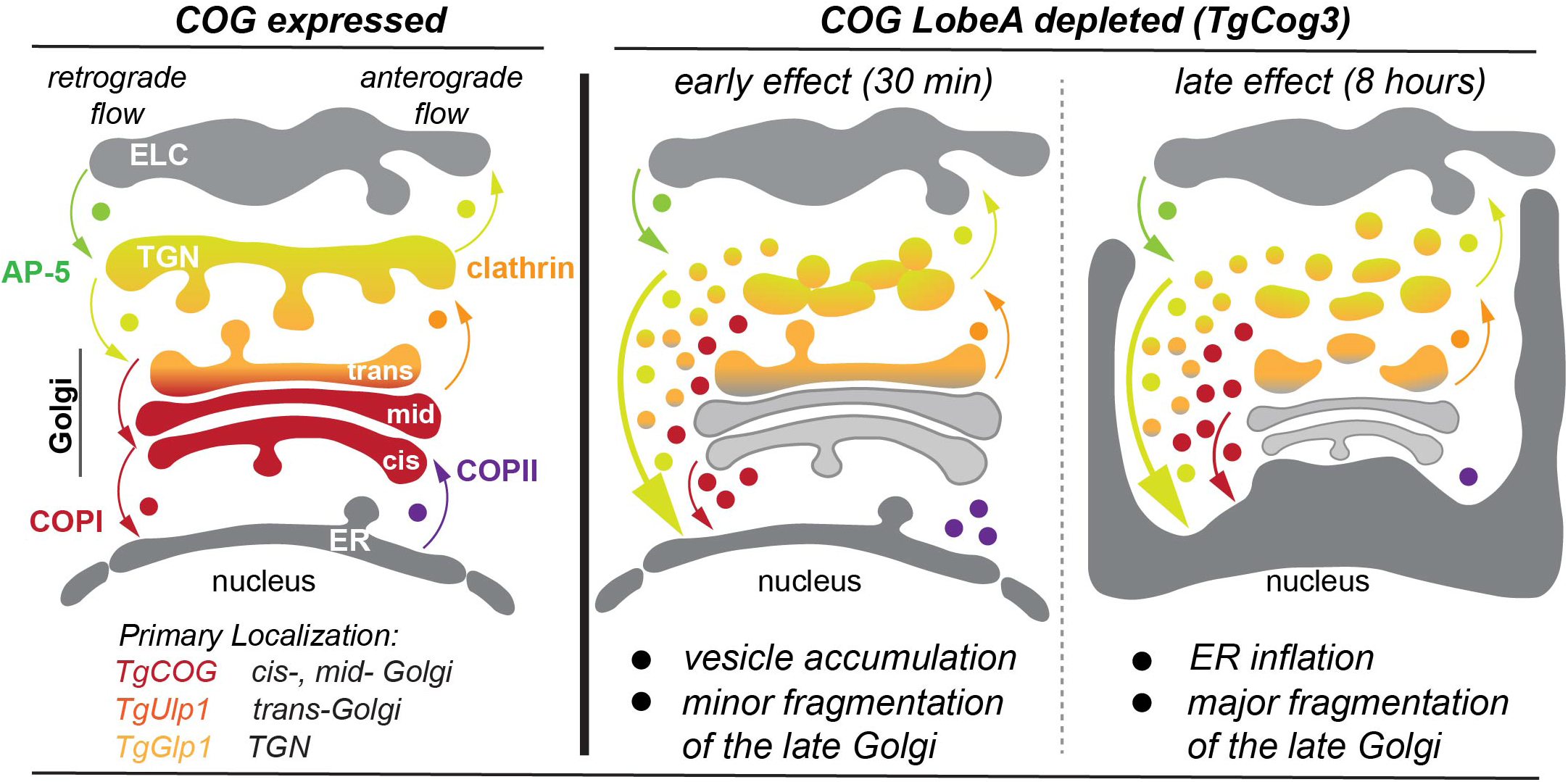
The model of the COG complex function in *T. gondii*. The drawing depicts a portion of the secretory pathway (from bottom to top): endoplasmic reticulum (ER); cis-, mid- and trans-Golgi; Trans Golgi Network (TGN); and endosome-like compartment (ELC). The organelles are colored according to the predicted localization of the COG complex (red), TgUlp1 (orange) and TgGlp1 (yellow). The arrows show direction of the vesicular transport and the type of transported vesicles. The lobe A subunit TgCog3 predicted to localize on acceptor Golgi membrane. The immediate effect of the acute TgCog3 degradation (30 min) is the block of the Golgi receiving function. In the absence of the Golgi tether, the anterograde and the retrograde vesicles cannot dock or fuse with Golgi and accumulate in the cytoplasm. The large membranous compartment with functional tethers, ER, likely accepts the rogue vesicles. Continued TgCog3 deprivation (8 hours) leads to ER inflation and impaired anterograde transport contributes to the phenotype. The created disbalance of the anterograde and retrograde flows results in Golgi reduction and fragmentation of the late Golgi compartments.

We made an intriguing discovery that suggests *T. gondii* has reinvented missing components of protein transport machinery. Two novel coccidian specific factors detected in the TgCog8 pulldown appear to be a new addition to retrograde Golgi transport. The COG complex interactor TgUlp1 displays structural features of the missing coiled coil Golgi tether Uso1/p115 [59]. Despite that, TgUlp1 role is substantially different from that of Uso1, which support the hypothesis that this factor was adapted rather than inherited. The function of conventional Uso1/p115 is limited to the ER-Golgi region where it interacts with COPII and COPI vesicles, whereas our analyses placed *T. gondii* Uso1-like factor in the late Golgi compartment (trans-Golgi or TGN) where it receives recycling AP-5 vesicles arriving from endosomal system, and to some extent, the recycling intra-Golgi COPI vesicles (Fig. 8). Like TgUlp1, another factor identified in this study TgGlp1 is a coccidian-specific protein with no distinctive structural features besides the presence of short, coiled coil regions in its C-terminus. Although the TgGlp1 and TgUlp1 do not interact with one another, their interactomes are nearly identical, indicating that TgUlp1 and TgGlp1 are involved in the same pathway. TgGlp1 and TgUlp1 are long proteins with coiled coil domains that are often present in molecular tethers. Both factors favor interactions with retrograde transport machinery (AP-5 and COPI vesicles) and have an identical set of Golgi and endosomal SNAREs. However, we detected a significant difference that places TgGlp1 and TgUlp1 in different Golgi regions. Surprisingly, downregulating the COG complex affected localization of TgGlp1 and not the direct COG complex partner TgUlp1. Upon downregulation of the lobe A subunit TgCog3, TgGlp1 was quickly redistributed to Golgi associated vesicles, while TgUlp1 remained associated with the Golgi. This is reminiscent of the differential reaction of Golgi coiled coil tethers to COG dysfunction in human cells. While Uso1/p115 demonstrated COG independent Golgi localization, other tethers like GOLGB1/giantin and GOLGA5/golgin-84 were quickly redistributed to CCD vesicles [29]. One plausible explanation is that TgGlp1 associates with trans-Golgi or TGN that donates recycling vesicles to preceding Golgi compartments, which is regulated by COG complex. Thus, continuously budding TGN retrograde vesicles with attached TgGlp1 cannot fuse with acceptor Golgi membranes that lack major Golgi tethering COG complex. We could not confirm direct TgGlp1-COG interaction, making the alternative less likely, where COG complex would control TgGlp1 association with Golgi. Nevertheless, the substantial overlap in COG complex localization and function with TgUlp1 and TgGlp1 suggests that they are part of the same trafficking machinery.

Why has *T. gondii* evolved novel transport factors? We found a rational answer in the interactomes of novel factors that were enriched in over twenty IMC proteins and suture factors, suggesting TgGlp1 and TgUlp1 involvement in the biogenesis of the parasite-specific IMC compartment. Why are these factors present only in coccidian parasites? Perhaps since coccidian parasites replicate by internal budding, which requires recycling of IMC material, and in turn requires expanding and adapting retrograde transport. In support of our hypothesis, TgGlp1 and TgUlp1 interactomes also contained Rab11b, which was previously shown to regulate IMC trafficking [66]. Although we detected some rhoptries, microneme, and dense granule factors in TgGlp1 and TgUlp1 pulldowns, their number and abundance were not comparable to IMC factors. Thus, the important conclusion of our study is that *T. gondii* supplemented and broaden the function of the conservative transport machinery such as COG complex to accommodate the specific needs of the parasite. The novel components mimicking conventional machinery (Uso1/p115 like factor) may have evolved to integrate the service of novel organelles into a conserved membrane trafficking system.

## MATERIAL AND METHODS

### Parasite cell culture

*T. gondii* RH*ΔKu80Δhxgprt AtTIR1* and RH*TatiΔKu80* strains were maintained in Human foreskin fibroblasts (HFF) (ATCC, SCRC-1041) cells in DMEM media (Millipore Sigma). Mycoplasma Detection (MP Biomedicals) PCR kit was used to confirm that parasite strains and HFF cells were free of mycoplasma. The transgenic lines used in this study are listed in Table S1.

### Phylogenetic analysis

Protein sequences were downloaded from UniProt database and analyzed in NGPhylogeny (www.NGPhylogeny.fr.) [67, 68]. Custom workflow included MUSCLE algorithm and BMGE to align and curate sequences [69, 70]. Maximum likelihood-based inference with Smart Model Selection was implemented using PhyML+SMS algorithm [71]. We applied 1000 bootstraps to test the optimization of each edge of the tree. The resulting phylogenetic trees were created using Newick utilities [72]. Sequences of the analyzed portions are included in Table S2.

### Construction of transgenic strains

Table S1 lists transgenic lines and primers used. Targeting constructs were verified by sequencing and PCR using gene- and epitope tag-specific primers was used to ensure that the tags were incorporated properly in their genomic locus.

#### Endogenous C-terminal tagging and conditional expression

To create AID conditional expression models for TgCog2, TgCog4, TgCog5, TgCog6, TgCog7, and TgCog8, genomic fragments of the 3’-end of the gene of interest were amplified by PCR and cloned into the pLIC mAID_3xHA_HXGPRT vector digested with PacI endonuclease by Gibson assembly method. Resulting constructs were linearized within the cloned gene fragment and transfected into RH*ΔKu80Δhxgprt AtTIR1* parent. A similar approach was used to build 3xmyc-epitope tagged TgUlp1, TgGlp1, TgCOPI-δ, TgSec31, and TgCog3. Genomic fragments were cloned into pLIC 3xmyc_DHFR-TS vector prior to linearization and transfection.

#### Endogenous N-terminal tagging and conditional expression

To build tet-OFF mutants of TgCog1 and TgCog2, the 5’ ends of the genes were amplified (Table S1 lists primers used), digested with BglII/NotI (TgCog2) or BamHI/NotI (TgCog1), and ligated into the promoter replacement vector pTetO7sag4-3xHA_DHFR-TS [44]. The resulting TgCog1 and TgCog2 tet-OFF constructs were linearized and transfected into RH*TatiΔKu80* strain.

#### Endogenous C-terminal tagging by CRISPR and conditional expression

To introduce mAID-3xHA epitope the C-terminus of TgCog3, we used the CRISPR/Cas9 approach as previously described [39]. We created pSAG1:CAS9-GFP U6: gsTgCog3 plasmid by site-specific mutagenesis of pSAG1:CAS9-GFP U6: gsUPRT plasmid (generously provided by Dr. David Sibley) to introduce a double DNA break in the 3’UTR of *TgCog3* locus [73]. A tagging cassette containing the selection marker *HXGPRT* flanked by 40bp gene-specific sequences amplified by PCR was co transfected with pSAG1:CAS9-GFP U6: gsTgCog3 plasmid into the RH*ΔKu80Δhxgprt AtTIR1* parent.

#### Parasite transfection and selection

Parent strain tachyzoites were grown in HFF monolayer. Freshly lysing parasites were collected and mixed with 50 µg DNA in 100 µl Cytomix buffer supplemented with 20mM ATP and 50mM reduced glutathione. Electroporated parasites (Amaxa, Lonza) underwent a 24-hour recovery in regular growth medium prior to drug selection. Drug-resistant polyclonal populations were cloned by limiting dilution. The resulting clonal population was screened using immunofluorescence microscopy with antibodies against the tagged protein of interest. Recombination at the target locus was confirmed by PCR amplification. Expression of tagged proteins was verified by Western blot analysis.

### Growth analysis

#### Plaque Assay

Plaque assays were performed in 6-well plates. Confluent HFF monolayers were infected with 50 parasites per well and treated with 500 μM Indole-3-Acetic Acid (IAA, auxin) which prompts degradation of mAID tagged proteins. Viability of tet-OFF parasites was evaluated by treating infected monolayers with 1μg/ml anhydrotetracycline (ATc). Plaques developed for 7 days at 37°C and then stained with crystal violet and counted. Three biological replicates of each assay were performed.

#### Vacuole size

The number of parasites per vacuole was determined after 16h growth at 37°C. Quantifications were done on 50-100 randomly selected vacuoles.

#### Immunofluorescent Microscopy Analysis

HFF infected parasites were grown on glass coverslips, fixed, permeabilized, and then incubated with the desired antibody. The following antibodies were used: rabbit a-Myc (clone 71D10; Cell Signaling Technology), rat a-HA (clone 3F10; Roche Applied Sciences), mouse a-Centrin (clone 20H5; Millipore Sigma), and rabbit a-IMC1 kindly provided by Dr. Gary Ward (University of Vermont, VT). A dilution of 1:500 was used for Alexa conjugated secondary Antibodies (Thermo Fisher Scientific). Nuclei were stained using 4’,6-diamidino-2-phenylindole (DAPI, Sigma). Glass cover slips were mounted in ImmunoMount (Thermo Fisher Scientific) and analyzed under a Zeiss Axiovert Microscope with Apotome optical slicer. Images were processed in Zen Blue 2.0 and Adobe Photoshop 2023. Single slice images were collected on a Zeiss LSM 880 inverted microscope with Airyscan and colocalization analysis was performed in the Zen Blue 2.6 colocalization module (Pearson coefficient) and 3D reconstructions were generated from confocal image Z-stacks.

### Transmission electron microscopy

The TEM Samples were processed according to published protocol with some modifications [74, 75]. Briefly, infected HFF monolayers were rinsed and fixed with 1% paraformaldehyde (PFA) (EMS) and 2.5% glutaraldehyde (GA) (EMS) in PBS for 2h at room temperature and stored in 1% PFA in PBS at 4°C and finally fixed with 2.5% GA and 0.05% malachite green (EMS) in 0.1 M sodium cacodylate buffer, pH 6.8 (20 min, ice). Then, cells were washed with 0.1M sodium cacodylate buffer, post-fixed in 0.5% osmium tetroxide and 0.8% potassium ferricyanide in 0.1 M sodium cacodylate buffer (30 min, room temperature), washed and incubated in 1% tannic acid (20 min, ice) and incubated in 1% uranyl acetate (1h, room temperature). Specimens were gradually dehydrated in increasing ethanol concentrations, washed with Propylene Oxide (EMS) and incubated in 50% PO/resin mixture before embedding in Araldite 502/Embed 812 resins (EMS). Ultrathin 50 nm sections were post-stained with aqueous uranyl acetate (EMS) and Reynold’s lead citrate (EMS). Images were acquired on FEI Technai G2 TF20 intermediate voltage transmission electron microscope at 80 keV (FEI Co.) equipped with FEI Eagle 4k CCD Camera and processed with FEI software.

### Immunoprecipitation assay

Protein samples were prepared from at least 2x10^9^ parasites collected by filtration and centrifugation. Total proteins were extracted in phosphate buffered saline solution with 263mM NaCl, 1% Triton X-100, and Halt^TM^ protease and phosphatase inhibitor cocktail (Thermo Scientific). Proteins were isolated on α-HA or α-Myc magnetic beads (Mbl Life Science). Pulldown efficiency was verified by western blot analysis.

### Western blot analysis

Protein samples made from isolated parasites or protein extracts were mixed with Laemmli sample buffer, heated at 95°C for 10 minutes, and sonicated. Proteins were separated via SDS PAGE and transferred onto Nitrocellulose membranes. Membranes were probed with the following primary antibodies: rat α-HA (clone 3F10; Roche Applied Sciences), rabbit α-Myc (Cell Signaling Technology), mouse α-GRA7 (kindly provided by Dr. Peter Bradley, UCLA CA), and mouse α-Tubulin A (12G10; kindly provided by Dr. Jacek Gaertig, University of Georgia, Athens GA). To visualize antigens, membranes were incubated with α-rat, α-mouse or α-rabbit secondary antibodies conjugated with horseradish peroxidase (Jackson ImmunoResearch). Proteins were visualized using enhanced chemiluminescent detection kit (Millipore). To detect changes in the protein glycosylation, membranes with separated proteins were blocked in Bio-Rad blocking buffer and probed with lectins as previously described [29]. Briefly, *Helix pomatia* Agglutinin (HPA) conjugated with Alexa-647 (Thermo Fisher Scientific) or biotinylated Concanavalin A and Jacalin (Vector) were diluted 1:1000 in Bio-Rad blocking buffer and incubated with membranes for 1 hour at room temperature. Biotinylated proteins were detected with 30 min additional incubation with Streptavidin-Alexa 647 (Thermo Fisher Scientific). Membranes were washed four times in PBS and imaged using the Odyssey Imaging System.

### Proteomic Analysis

#### Mass-spectrometry

Protein samples were prepared for mass spectrometry-based proteomic analysis using s-traps (Protifi) as previously described [76, 77]. Proteins were reduced with 20 μM dithiothreitol (DTT) for 10 minutes at 95°C, followed by a 30-min alkylation with 40 μM Iodoacetamide (IAA). Proteins were acidified using phosphoric acid and combined with s-trap loading buffer (90% methanol, 100 mM triethylammonium bicarbonate). The resulting modified proteins were loaded onto s-traps, washed, and digested with trypsin/Lys-C (1:100 wt/wt, enzyme/protein) overnight at 37°C. Peptides were eluted, dried in a vacuum concentrator, and then resuspended in H2O–1% acetonitrile–0.1% formic acid for liquid chromatography tandem mass spectrometry (LC-MS/MS) analysis. Peptides were separated using a 75µm x 50cm C18 reversed-phase-UHPLC column (Thermo Scientific) on an Ultimate 3000 UHPLC (Thermo Scientific) with a 120-minute gradient (2-32% acetonitrile with 0.1% formic acid). Full MS survey scans were acquired at 70,000 resolution. Data-dependent acquisition (DDA) selected the top 10 most abundant ions for MS/MS analysis. Raw data files were processed in MaxQuant (www.maxquant.org) and searched against the current ToxoDB (www.toxodb.org) *Toxoplasma gondii* ME49 protein sequence database. Search parameters include constant modification of cysteine by carbamidomethylation and the variable modification, methionine oxidation. Proteins were identified using 1% FDR (false discovery rate) filtering criteria. The data is available in the PRIDE proteomics exchange database (www.proteomexchange.org).

#### Data analysis

The Significance Analysis of INTeractome (SAINT) was used to identify TgCog8, TgUlp1 and TgGlp1 interactors with default parameter setting in function ‘SAINT express version’. We incorporated proteomes of the parental strains RH*ΔKu80Δhxgprt AtTIR1* and RH*ΔKu80* to account for non-specific binding.

### Structural analysis of hypothetical proteins

To identify homology models, amino acid sequences of the target proteins were examined in the SWISS-MODEL suite [78]. The resulting models were further analyzed in PyMol (www.pymol.org). Available images of the folded proteins were downloaded from AlphaFold2. Alignment of the protein sequences downloaded from UniProt (www.uniport.org) database was performed using MUSCLE algorithm and exported to Jalview [79]. The domain of interest within each protein was selected and displayed using Clustal color scheme.

## SUPPLEMENTAL FIGURE LEGENDS

**Figure S1.**
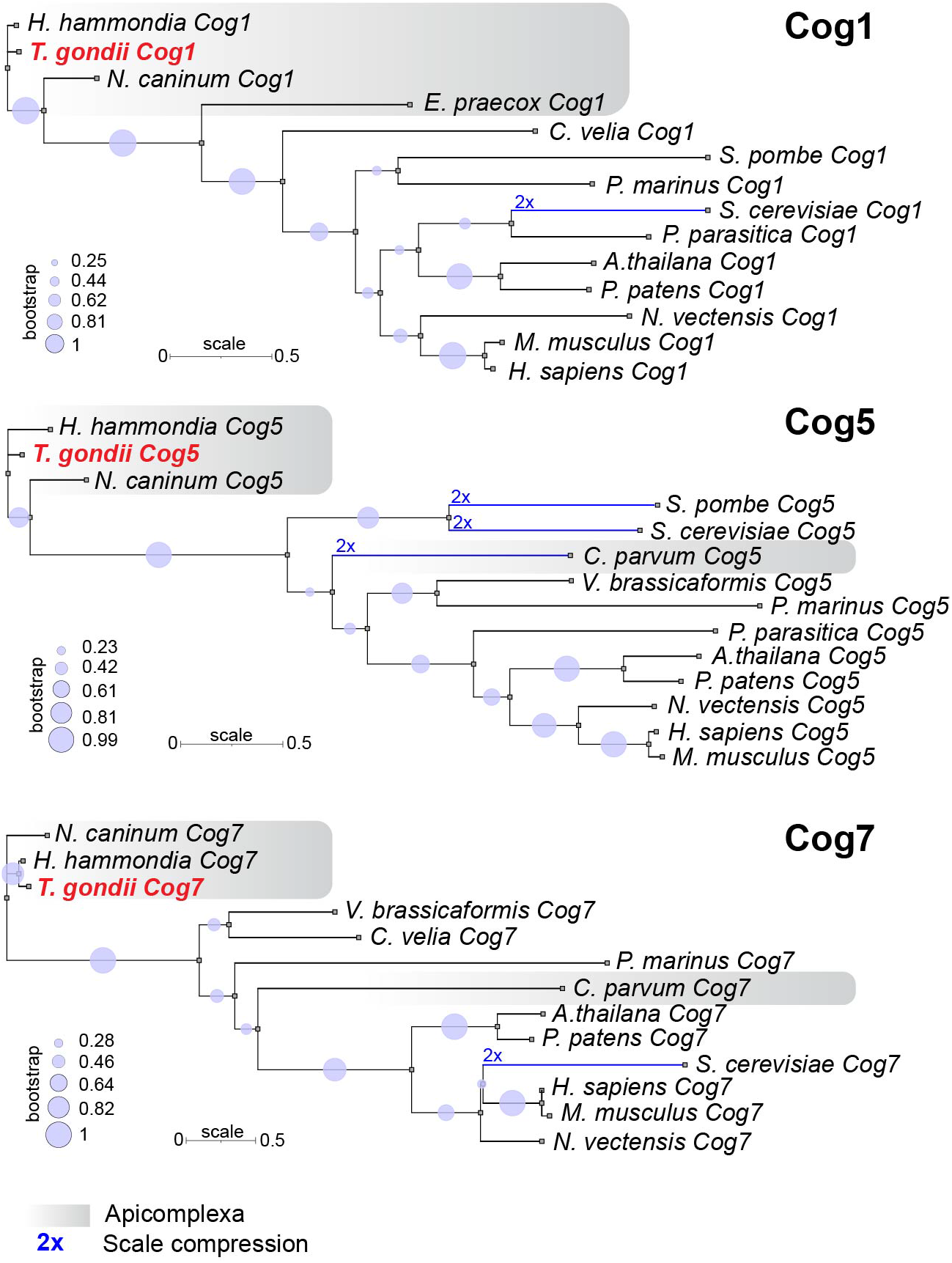
Phylogenetic analysis of Toxoplasma COG complex subunits TgCog1, TgCog5 and TgCog7. Shaded blocks show apicomplexan orthologs. Blue line is a compressed tree branch, the compression coefficient is indicated.

**Figure S2.**
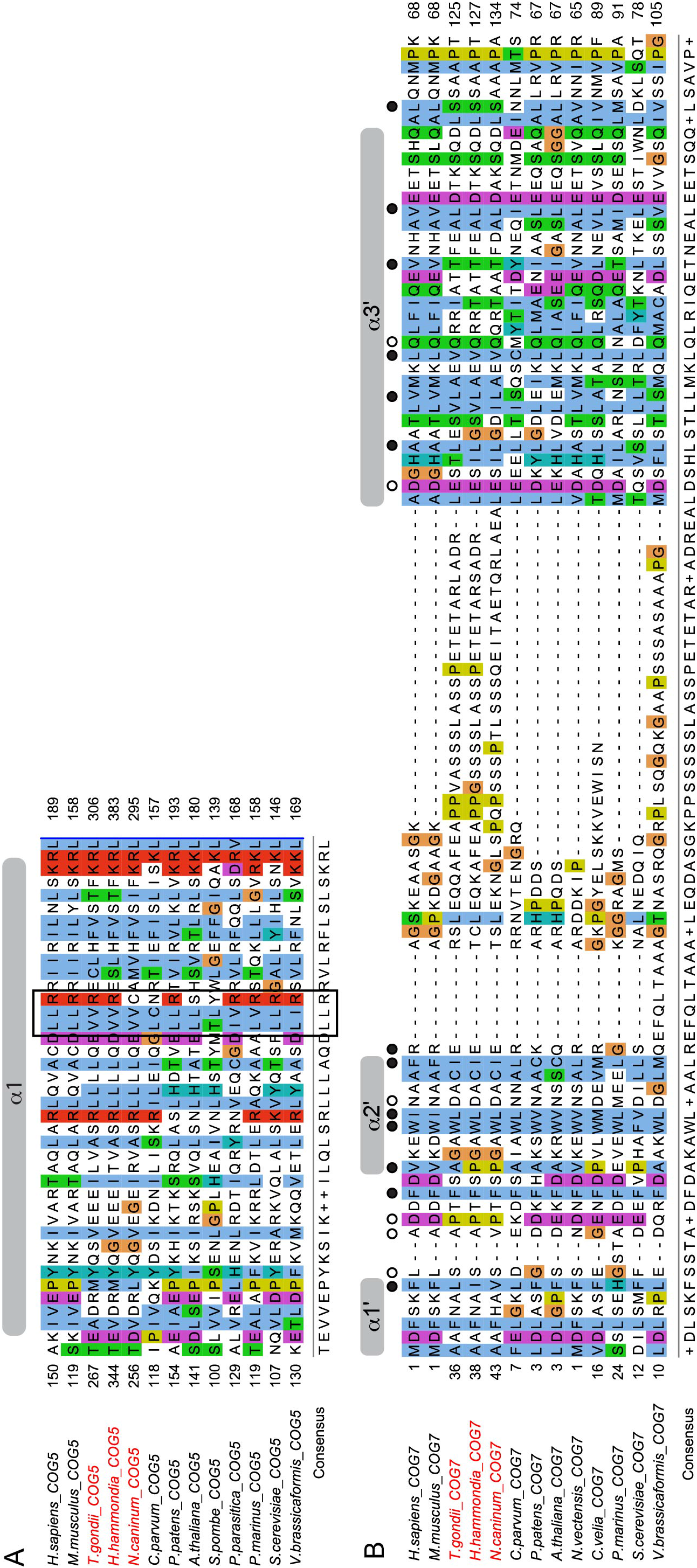
The TgCog5 and TgCog7 signature domains. **A.** Alignment of the COG5 CATCHR domain (MUSCLE). Secondary structure (above) and sequence conservation (below) are shown, including central LLR (VVR) motif. **B.** Alignment of the COG7 α1’, α2’, and α3’ helixes that are involved in interaction with COG5 CATCHR domain (MUSCLE). Critical amino acid residues are marked with closed (non-polar interaction) and open (polar interaction) circles.

**Figure S3.**
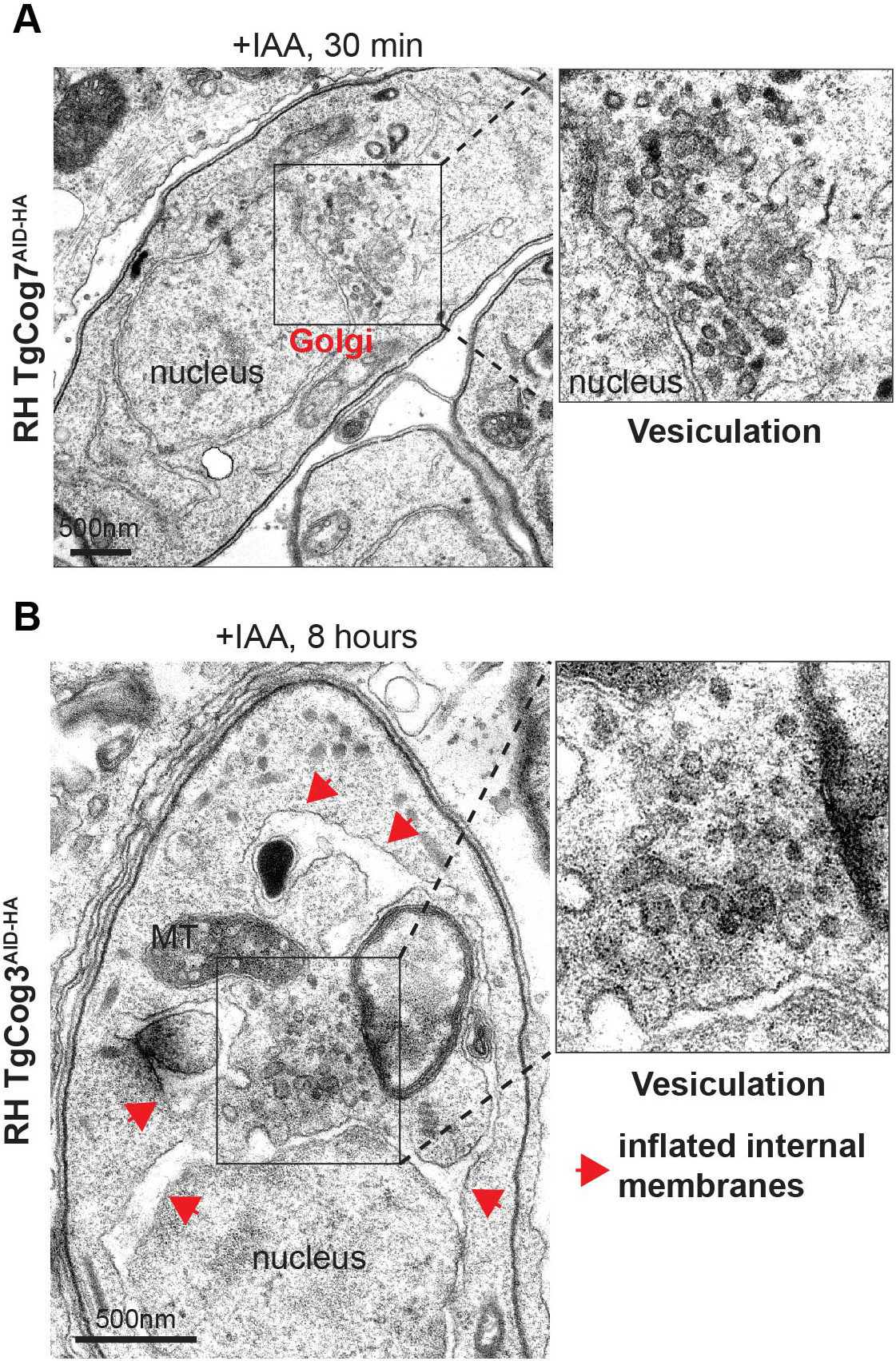
TEM analysis of *T. gondii* COG complex deficiency. Images of TgCog7 (A) and TgCog3 (B) depleted parasites show persisted vesiculation in the Golgi region.

**Figure S4.**
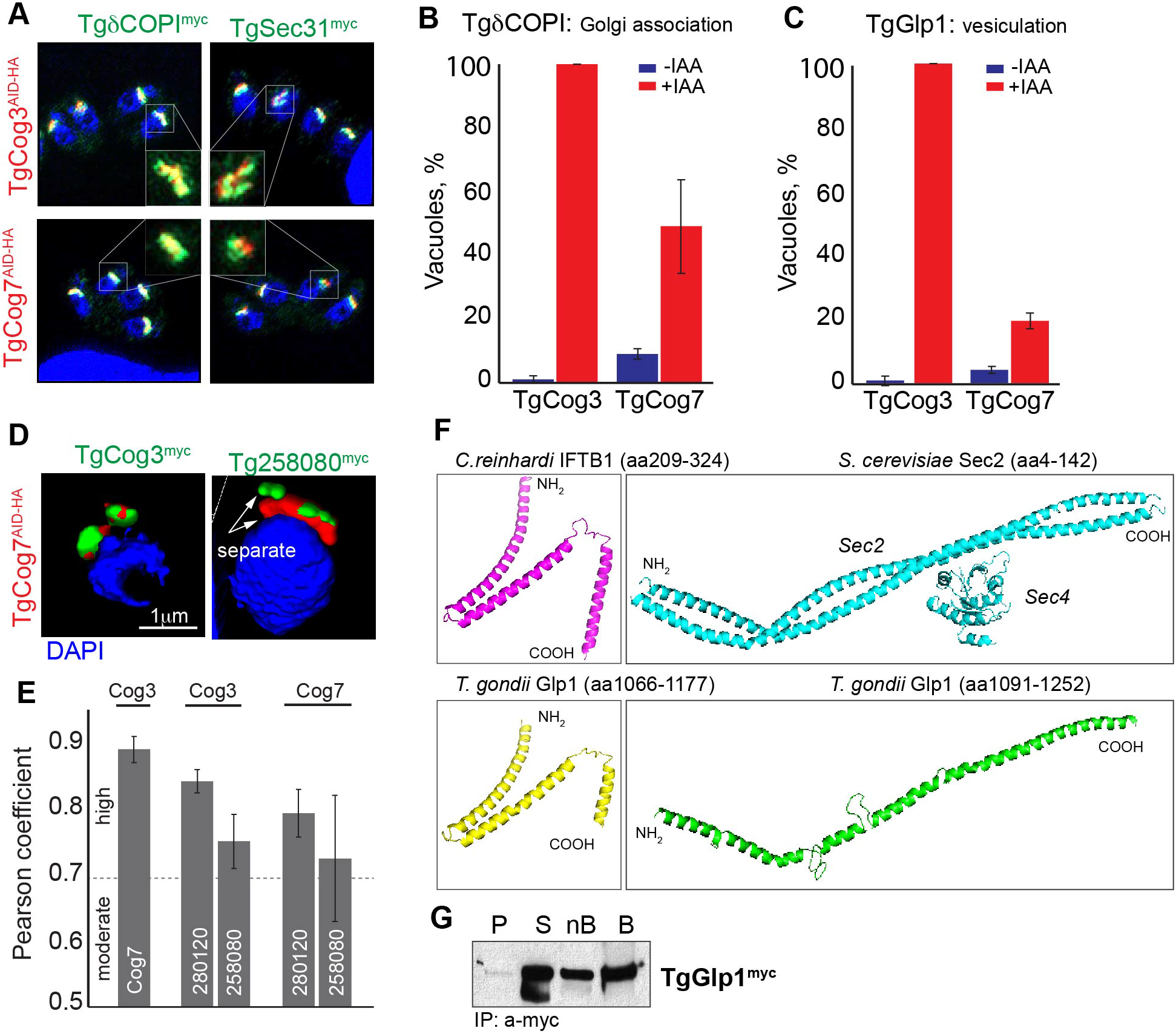
Analysis of the *T. gondii* COG complex interactions. A. Immunofluorescent analysis of the parasites expressing TgδCOPI^myc^ or TgSec31^myc^ in the RH TgCog3^AID-HA^ or TgCog7^AID-HA^ mutants. Parasites were co-stained with α-myc (green), α-HA (red) and DAPI (blue). Insets shows the proteins overlap in the Golgi region. **B.** Quantification of the TgδCOPI Golgi association in parasites not treated or treated with IAA for 8 hours. The mean of the vacuole counts, and the SD values are plotted on the graph. **C.** Quantification of the parasites that showed Golgi vesiculation (TgGlp1^myc^ marker) upon 8-hour treatment with IAA. The mean of the vacuole counts, and the SD values are plotted on the graph. **D.** A 3-D reconstruction of the Immunofluorescent images of the parasites expressing TgCog3^myc^ or Tg258080^myc^ in the RH TgCog7^AID-HA^ line. Parasites were co-stained with α-myc (green), α-HA (red) and DAPI (blue). **E.** Pearson coefficients of TGME49_258080, TGME49_289120 proteins colocalization with the COG complex subunits TgCog3 and TgCog7. The mean and the SD values collected from a minimum 10 parasites are plotted on the bar graph. **F.** Folding prediction for selected regions of *T. gondii* Glp1 (bottom panel) and models (top panel) are shown (PyMol). Note that *S. cerevisiae* Sec2 shown in the complex with Sec4. **G.** Western Blot analysis of immunoprecipitated TgGlp1^myc^ complexes. The insoluble [P, pellet], soluble [S], depleted soluble fractions [nB, not bound] and the beads with precipitated complexes [B] (10 times more than the other fractions) were probed with α-myc antibodies.

